# Genome Sequences of *Burkholderia thailandensis strains E421, E426, and DW503*

**DOI:** 10.1101/2020.01.24.915645

**Authors:** Catherine M. Mageeney, Anupama Sinha, Kelly P. Williams, Steven S. Branda

## Abstract

We present the draft genome sequences of three *Burkholderia thailandensis* strains: E421, E426, and DW503. E421 consists of 90 contigs of 6,639,935bp and 67.73% GC content. E426 consists of 106 contigs of 6,587,853bp and 67.73% GC content. DW503 consists of 102 contigs of 6,458,767bp and 67.64% GC content.

## Main text

*Burkholderia thailandensis* is a Gram-negative bacterium that occurs naturally in soil and water. It is commonly used as a surrogate for study of its more pathogenic relatives, the Select Agents *Burkholderia pseudomallei* and *B. mallei*, with which it shares many genome features and pathogenesis mechanisms, and provokes similar immune responses in mammalian systems (1). Here we report the genome sequences of three *B. thailandensis* strains obtained from the BEI Resources repository (Manassas, VA USA). *B. thailandensis* strains E421 and E426 (referred to as E421 and E426, respectively) were isolated independently from a rice field in Northeast Thailand; and *B. thailandensis* strain DW503 (referred to as DW503) is an allelic exchange mutant of *B. thailandensis* strain E264 (referred to as E264) in which the *amrR-oprA* operon is deleted (2).

E421, E426, and DW503 were each grown to confluency in monoculture, and their DNA extracted using the DNeasy Blood and Tissue Kit (QIAgen; Hilden, Germany). DNA sequencing libraries were generated using the Nextera DNA Library Prep Kit and the Nextera DNA Sample Preparation Index Kit (Illumina; San Diego, CA USA). Library prep concentrations were quantified using the Qubit fluorometer with its High Sensitivity DNA Assay Kit (Thermo Fisher Scientific; Waltham, MA USA). Equal amounts of the three individual libraries were mixed together to generate a multiplexed library, which was analyzed for concentration using the Qubit and for fragment size using the Bioanalyzer electrophoresis system with its High Sensitivity DNA Analysis Kit (Agilent; Santa Clara, CA USA). The multiplexed library was sequenced in paired-end mode using a NextSeq 500 with its High-Output (300 Cycles) Kit (Illumina).

Assembly of sequencing reads was completed using SPAdes (3) (Table 1). FastANI alignment (4) showed that E421 and E426 share 99.8% average nucleotide identity (ANI) with E264. As might be expected, DW503 shares 99.9978% ANI with E264. Sequence annotation was performed using our inhouse annotation pipeline, which calls Prokka (5), tFind, and rFind softwares (6); counts for coding regions, rRNA sequences, and tRNA sequences are listed in Table 1. All three strains had hits to the following Rfam (7) miscellaneous RNA profiles: 6S, Bacteria_large_SRP, Bacteria_small_SRP, Betaproteobacteria_toxic_sRNA, cspA, FMN, Intron_gpI, mini-ykkC, P9, pfl, RNaseP_bact_a, SAH_riboswitch, SECIS_3, sucA, and TPP; in addition, E421 and E426 hit the following: C4, Cobalamin, and isrK. Additionally, the sequences were analyzed to identify CRISPR-Cas arrays [CRISPR-CAS finder (8)], genomic islands [TIGER (Mageeney *et al*., submitted for publication) and Islander (6)], and, among the latter, prophages [in-house software (Mageeney *et al*., submitted for publication)]; Table 1 summarizes these data.

**Table 1.**
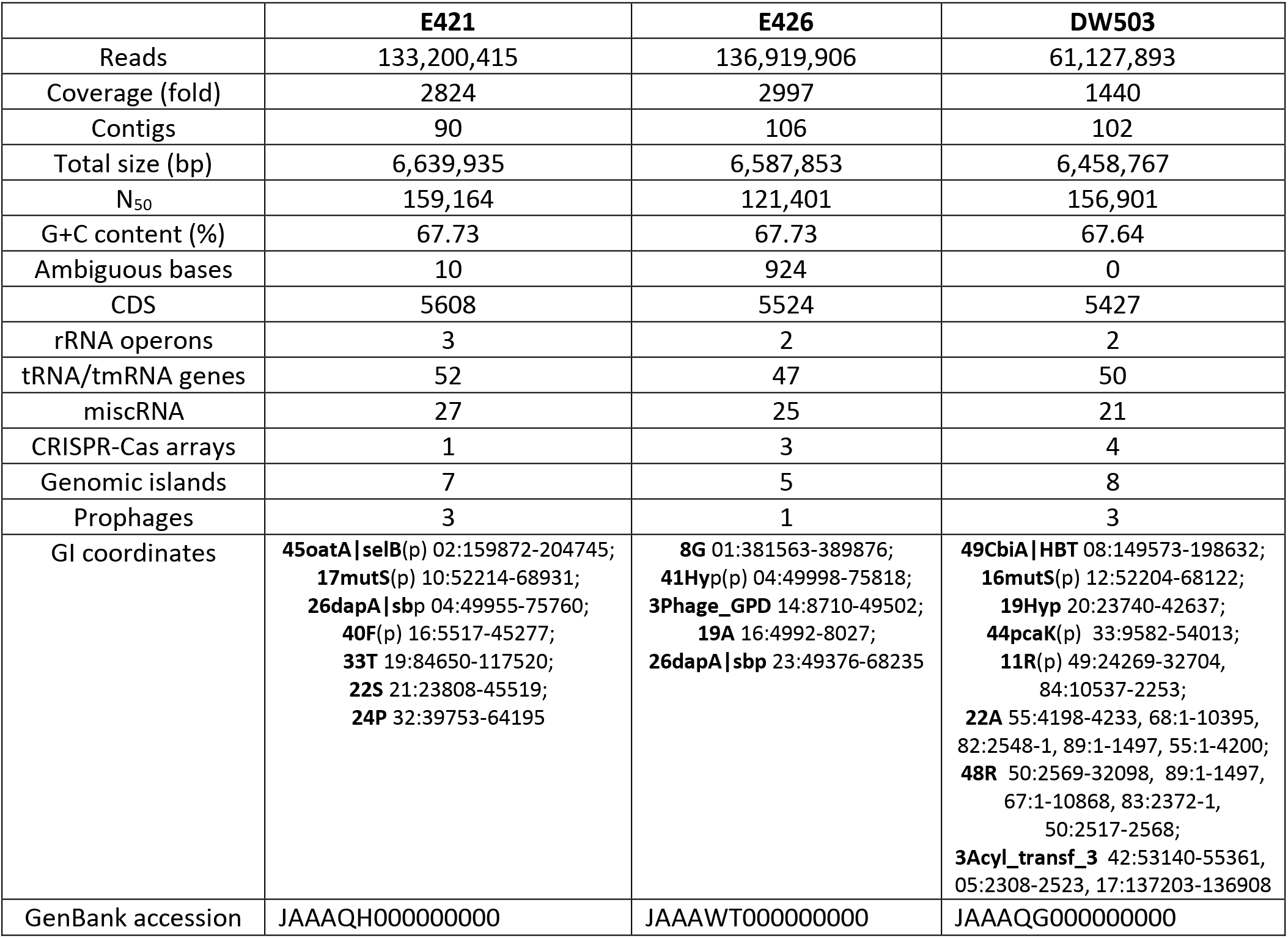
Assembly statistics and genome sequence features. Names (bold) of genomic islands (GIs), and coordinates with accession number, are listed; (p) indicates a prophage.

## Data availability

The sequencing data have been deposited at GenBank under the following accession numbers: JAAAQH000000000 (E421), JAAAWT000000000 (E426) and JAAAQG000000000 (DW503). The versions described in this paper are the first versions.

## Acknowledgements

Sandia National Laboratories is a multi-mission laboratory managed and operated by National Technology and Engineering Solutions of Sandia, LLC, a wholly owned subsidiary of Honeywell International, Inc., for the U.S. Department of Energy’s National Nuclear Security Administration under contract DE-NA0003525. E421 (NR-9909), E426 (NR-9908), and DW503 (NR-4075) were obtained through BEI Resources, NIAID, NIH. This paper describes objective technical results and analysis. Any subjective views or opinions that might be expressed in the paper do not necessarily represent the views of the U.S. Department of Energy or the United States Government.

